# Transcriptional response in a sepsis mouse model reflects transcriptional response in sepsis patients

**DOI:** 10.1101/2021.12.09.471887

**Authors:** Florian Rosier, Nicolas Fernandez Nuñez, Magali Torres, Béatrice Loriod, Pascal Rihet, Lydie C. Pradel

**Affiliations:** TAGC, Inserm UMR1090 Aix-Marseille Université, 163 avenue de Luminy, 13288 Marseille, France, 13288 Marseille, France; TGML-TAGC, Inserm UMR1090 Aix-Marseille Université, 163 avenue de Luminy, 13288 Marseille, France

**Keywords:** sepsis, mouse models, patients, Lipopolysaccharide, Microarray, monocytes/macrophages

## Abstract

Mortality due to sepsis remains unacceptably high, especially for septic shock patients. Murine models have been used to better understand pathophysiology mechanisms. However, the mouse model is still under debate. Here we investigated the transcriptional response of mice injected with lipopolysaccharide (LPS) and compared it to either human cells stimulated *in vitro* with LPS or to blood cells of septic patients. We identified a molecular signature composed of 2331 genes with an FDR median of 0%. This molecular signature is highly enriched in regulated genes in peritoneal macrophages stimulated with LPS. There is a significant enrichment in several inflammatory signaling pathways, and in disease terms, such as pneumonia, sepsis, systemic inflammatory response syndrome, severe sepsis, an inflammatory disorder, immune suppression, and septic shock. A significant overlap between the genes up-regulated in mouse and human cells stimulated with LPS has been demonstrated. Finally, genes up-regulated in mouse cells stimulated with LPS are enriched in genes up-regulated in human cells stimulated in vitro and in septic patients, who are at high risk of death. Our results support the hypothesis of common molecular and cellular mechanisms between mouse and human sepsis.

## 1. Introduction

Sepsis is a multifactorial clinical disease characterized by a dynamic clinical course and a diverse clinical outcome depending on several factors such as the physiopathology process, the environment and the phenotype of the affected patients. Sepsis results from a complex disruption of inflammation leading to an inability of the host to control an infection. Then, an exacerbated response of the immune system injures its own tissue and induces severe sepsis (sepsis with organ dysfunction) and septic shock, the worst complications of this pathology. It has been proposed more recently that an immunosuppression phase occurs later, leading to an increased risk of mortality. According to the World Health Organization (WHO), sepsis affects between 47 and 50 million people per year leading to 11 million deaths ([1] WHO Report on the burden of endemic health care-associated infection worldwide. 2017-11-21 15:11:22 2011). The prevalence of the disease is higher in low and middle-income countries. Sepsis can occur after trauma or any infections, including common bacterial illness but also Malaria or Covid 19. Murine models remain very useful to decipher the main causes of sepsis [2]. Among the available models, direct injection of toxins such as LPS into the blood, peritoneum or lung induces a strong inflammatory response that mimics the innate immune system in humans [3, 4]. The advantage of this method allows technical ease and a reproducible response of the immune system with the injection of a controlled amount of toxin. Here we performed a transcriptomic study from cells extracted from the peritoneal cavity after LPS injected in mice. We provide evidence that the mouse transcriptional response reflects both the response of human blood cells to bacteria and the risk of mortality in sepsis patients.

## 2. Results

### 2.1. Microarray gene expression profiling

Experimental design and data analysis were carried out as shown in Figure 1. In order to identify genes modulated by LPS injection, we isolated the peritoneal cells from 8 mice during the inflammatory peak 1 hour after the injection (n=4) and, just before the death, 40 hours after LPS injection (n=4), and from 4 control mice that were not injected with LPS. The RNA samples were further extracted and used for microarray gene expression profiling and qPCR experiments.

**Figure 1:**
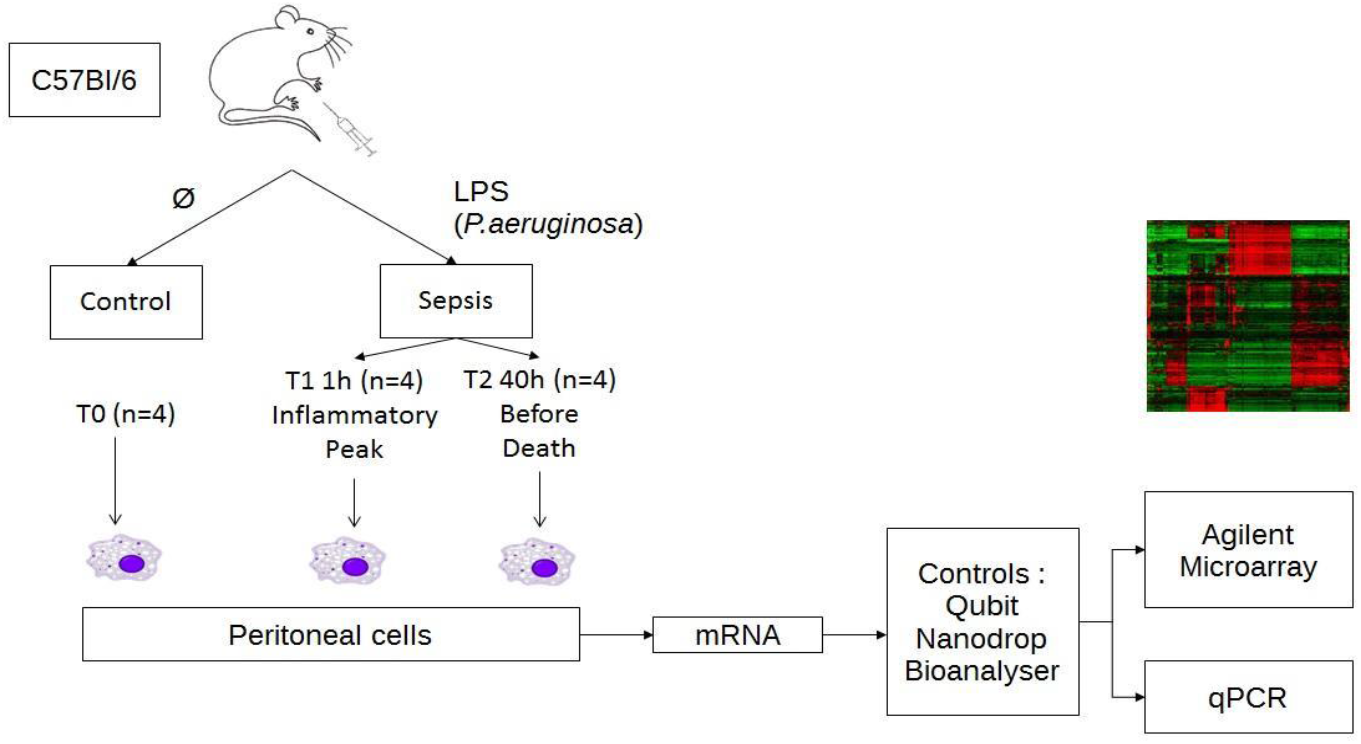
Experimental design and data analysis. Schematic outline representing the experimental design for the microarray analysis in SurePrint G3 8×60K mouse from Agilent. RNA was extracted from peritoneal cells of 4 mice (n=4 for each group) injected in peritoneal cavity either with PBS or with LPS during the inflammatory peak (T1=1h) and before death (T2=40h).

As many as 62,976 probes corresponding to 39,430 coding RNAs and 16,251 LincRNAs were used to detect mRNA expression levels for each RNA sample using SurePrint G3 Mouse Gene Expression 8 × 60 K v2 Microarray Kit, and the data were analyzed using “AgiND” library loaded into “R” for standardization. Probes with intensity values below the background noise were filtered out, resulting in 25,880 probes, which were used for differential expression analysis. Out of the 39,430 coding RNAs and 16,251 LincRNAs, the expression of 3169 probes were found to be statistically significantly different after Significance Analysis of Microarrays test with an FDR median of 0%.

Thus, 3,169 probes that were identified represented 2,361 genes differentially expressed in sepsis, including 130 LincRNAs (Supplementary Table 1A). After performing our hierarchical clustering (Figure 2), we observed that all our samples were correctly classified. In addition, samples from LPS-injected mice (T1 and T2) were observed to be closer than those of control mice (T0). We observed a clustering of our probes, which were separated into 6 clusters after removing the duplicated probes for genes and genes that clustered in several groups. Two thousand three hundred and thirty-one genes were still differentially expressed (Supplementary Table 1B). There were 109 genes in the first gene cluster, 1050 genes in cluster 2, 181 genes in the third cluster, 107 genes in the fourth cluster, 793 genes in the fifth cluster group genes, and finally 121 genes in cluster 6 (Figure 3).

**Figure 2.**
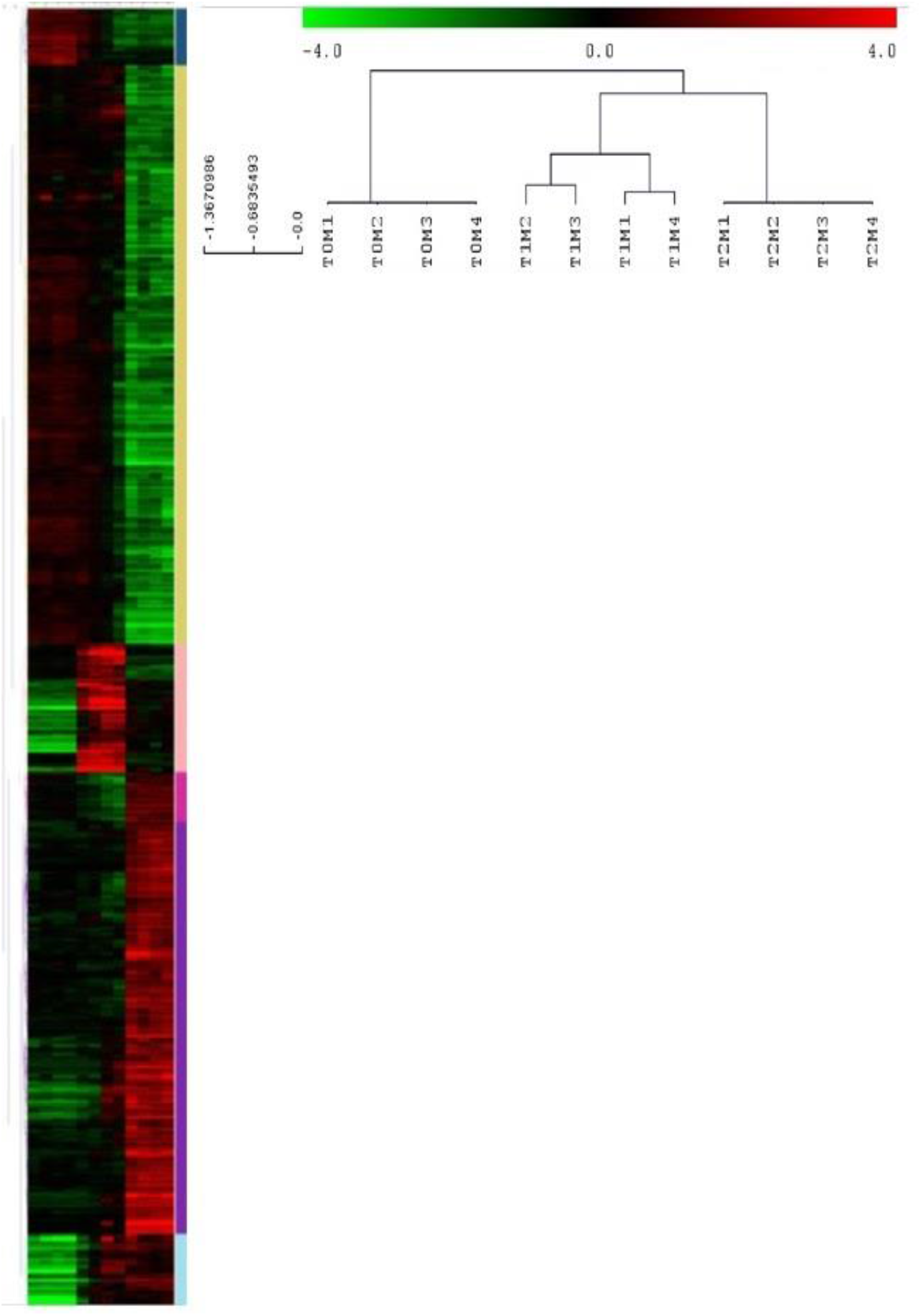
Unsupervised hierarchical clustering of probes in our mouse sepsis model. The samples in each column (vertical) and the probes in each row (horizontal) were (re)arranged. Six vertical clusters (blue, yellow, light pink, fuchsia, purple and light blue) and 3 horizontal clusters were obtained corresponding to genes of similar expression pattern placed close to each other, and samples with comparable traits (T0, T1 and T2), respectively. The heatmap indicates up-regulation (red), down-regulation (green), and mean gene expression (black).

**Figure 3:**
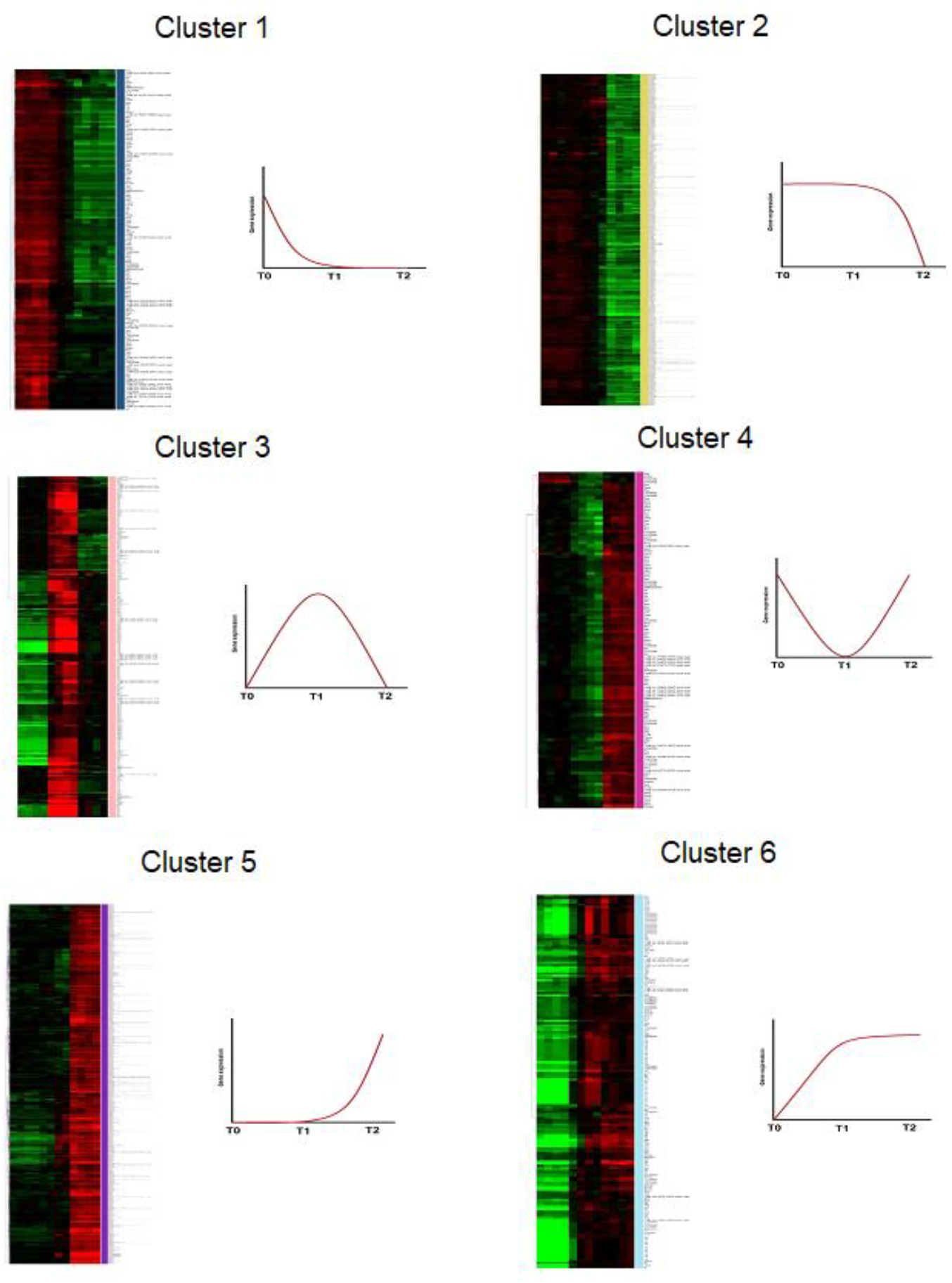
Hierarchical clustering of probes and schematic gene expression profile for each cluster identified in our mouse sepsis model (blue, yellow, light pink, fuchsia, purple and light blue). The genes expression profile corresponding to each heatmap cluster is shown to its right.

We further assessed the expression of 4 selected genes (TNF, IL1b, IL6, and IL10) by RT-qPCR to confirm the expression pattern. Gene expression level measured with microarray and with RT-qPCR were correlated for TNF (Spearman’s rho=0.882, P<0.001), IL1b (Spearman’s rho=0.891, P<0.001), IL6 (Spearman’s rho=0.845, P=0.002), and IL10 (Spearman’s rho=0.982, P<0.001).

Figure 4 shows differences in gene expression level. It highlights that gene expression patterns using microarray and RT-qPCR methods were consistent. In particular, statistical analyses of RT-qPCR measurements confirmed that there was a higher level of gene expression at T1 for the 4 genes (P<0.012).

**Figure 4:**
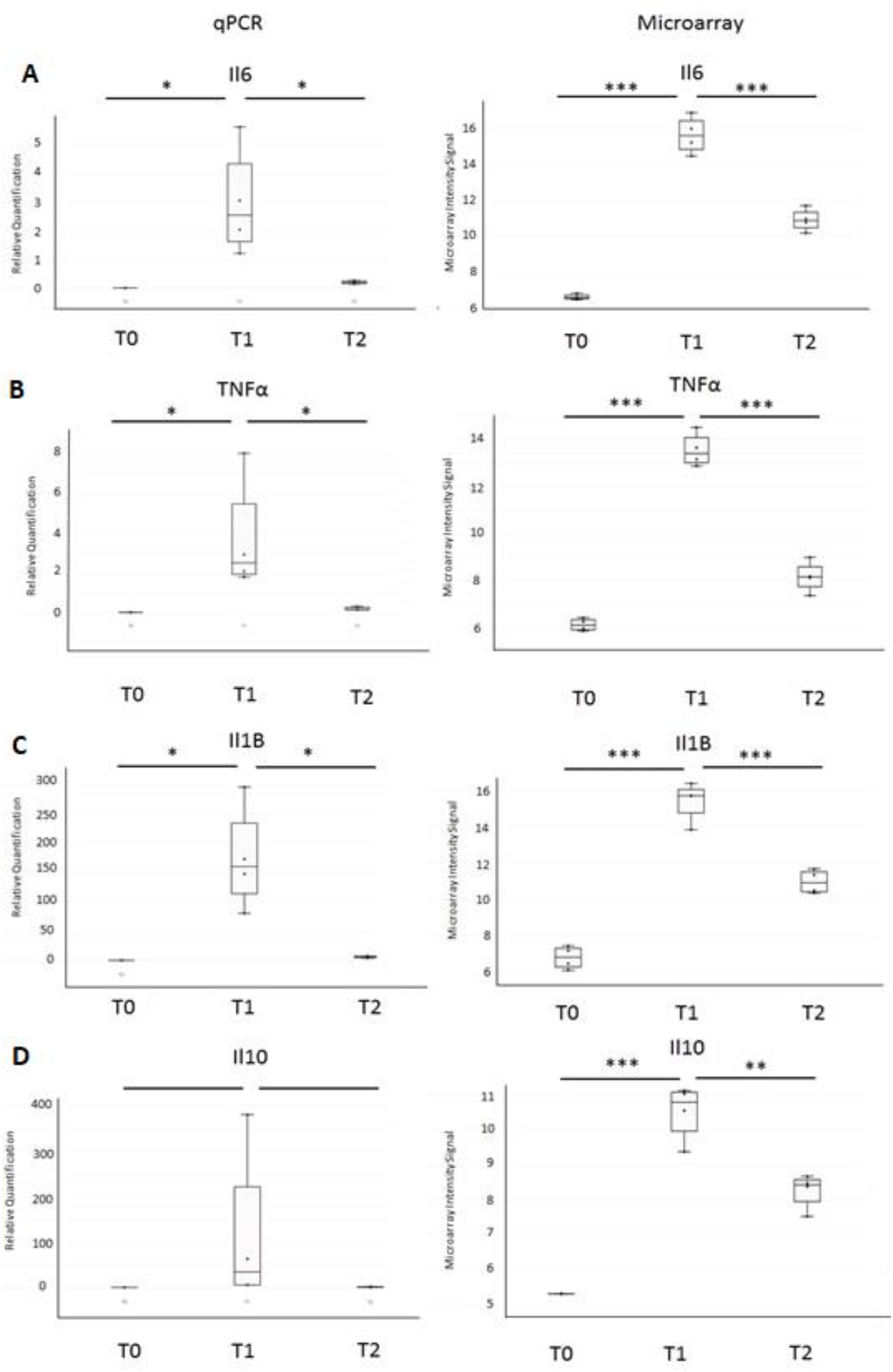
Expression levels of IL6 (A), TNFα (B), IL1b (C), and IL10 (D) measured by RT-qPCR (Left) and with microarray (right). For RT-qPCR, values were normalized with the actin transcript.

### 2.2. Functional microarray analysis

To analyze functional annotations related to mouse sepsis, we sought biological process Gene Ontology (GO) terms, mouse gene atlas terms, KEGG pathways, and disease ontology terms for the gene whose expression was up- or down-regulated in mice injected with LPS (Supplementary Table 2). The analysis of functional terms showed an over-representation of terms related to signaling pathways such as TNF, IL-17, cytokine-mediated, chemokine, NF-kappa B, Toll-like receptor, NOD-like receptor, MAPK, and JAK-STAT signaling pathways (Figure 5A).

**Figure 5:**
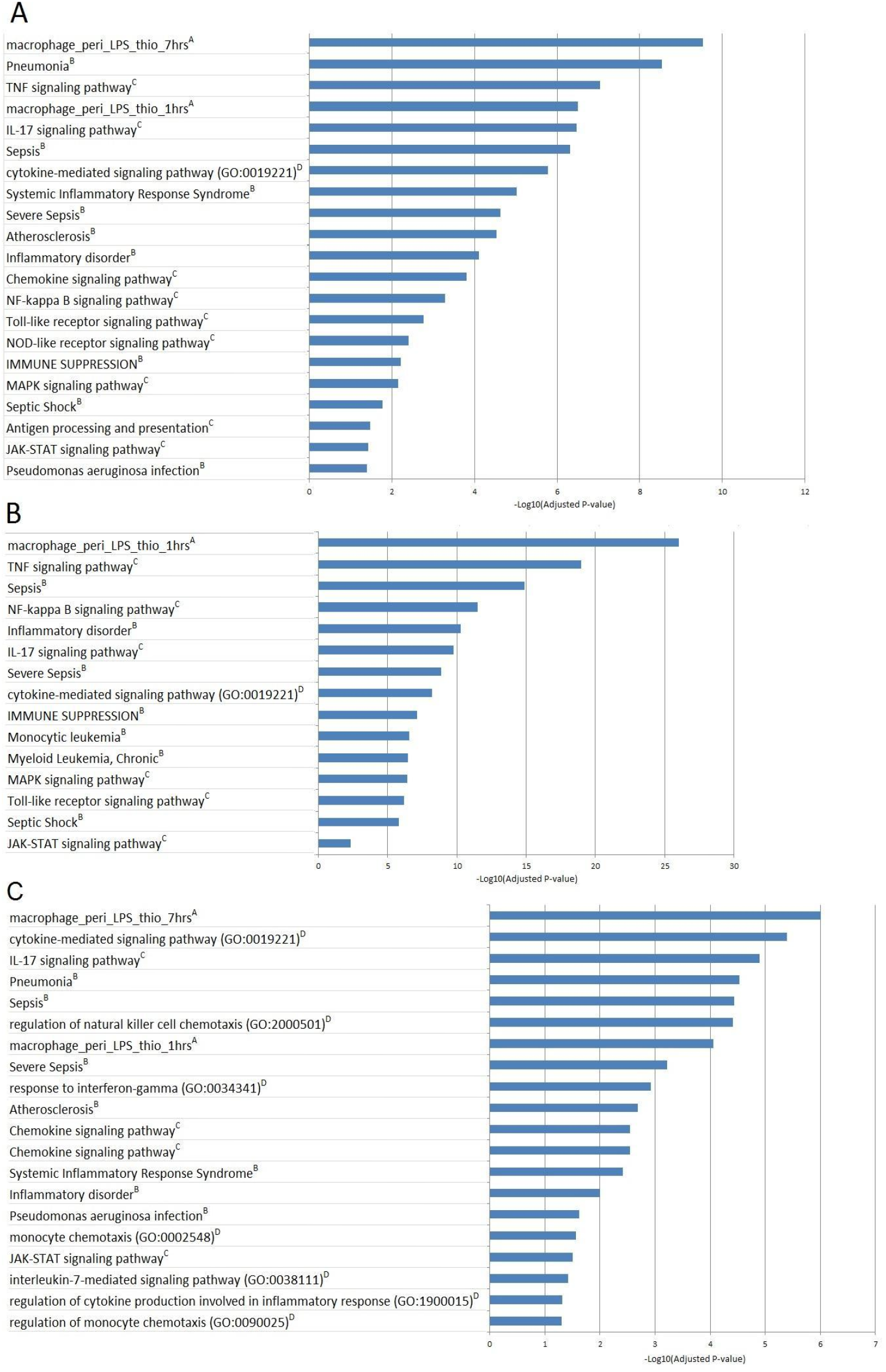
Biological processes identified by functional enrichment analysis for all clusters (A), cluster 3 (B) and cluster 6 (C). The negative Log10 of the adjusted pValue is represented.

Noticeably, the functional term that yielded the best enrichment P value was related to the response of peritoneal macrophages to LPS (Figure 5A,5B,5C). Furthermore, several disease ontology terms were significantly over-represented. These included pneumonia, sepsis, systemic inflammatory response syndrome, severe sepsis, inflammatory disorder, immune suppression, and septic shock. Most of those functional or disease terms were found when analyzing the enrichment in individual clusters, as shown for cluster 3 and 6 in Figure 5B and 5C, respectively. Moreover, there was an enrichment in terms that were specific for some clusters (Figure 5B and 5C). Also, we found that genes annotated with biological process ontology terms, such as “regulation of natural killer cell chemotaxis”, “response to interferon-gamma”, interleukin-7 mediated signaling pathway”, “regulation of cytokine production involved in inflammatory response”, “and regulation of monocyte chemotaxis”, were over-represented in the cluster 6 while the TNF and NF-Kappa B signaling pathways were over-represented in cluster 3.

### 2.3. Comparison of published lists of human differentially expressed genes with mouse differentially expressed genes

Moreover, we compared the list of 2331 differentially expressed genes in mice injected with LPS with transcriptional signatures in humans found in three papers (supplementary Table 3 and Table 2). First, we compared those mouse genes with the genes regulated in primary human monocytes stimulated with LPS for 2 hours or 24 hours (Figure 6E, 6F) [5]. There was a significant overlap between the human gene list and the mouse gene list (supplementary Table 3). We used the GSEA approach to assess the significance of the overlap between human differentially expressed genes and mouse differentially expressed genes (supplementary table 3). This is a robust approach, which reduces the bias due to different statistical methods and different stringency of cut-offs. Figure 6 shows that there was a significant enrichment of up-regulated genes in human cells stimulated with LPS for 2 (Figure 6E) and 24 hours (Figure 6F), compared to our transcriptome data, whereas there was no enrichment of down-regulated human genes, compared to down-regulated genes in mouse cells stimulated with LPS (Figure 7 and supplementary Figure 1). The GSEA approach allowed us to identify the leading edge gene subsets that mainly accounted for the enrichment signals: 183 and 196 genes upregulated in human cells stimulated with LPS for 2 hours or 24 hours were leading edge genes (Supplementary Table 3), respectively. The two leading edge gene sets were enriched in gene ontology terms related to inflammation, such as “inflammatory response”. Interestingly, *IL1B, IL6, IL10* and *TNF* were in both gene sets.

**Figure 6:**
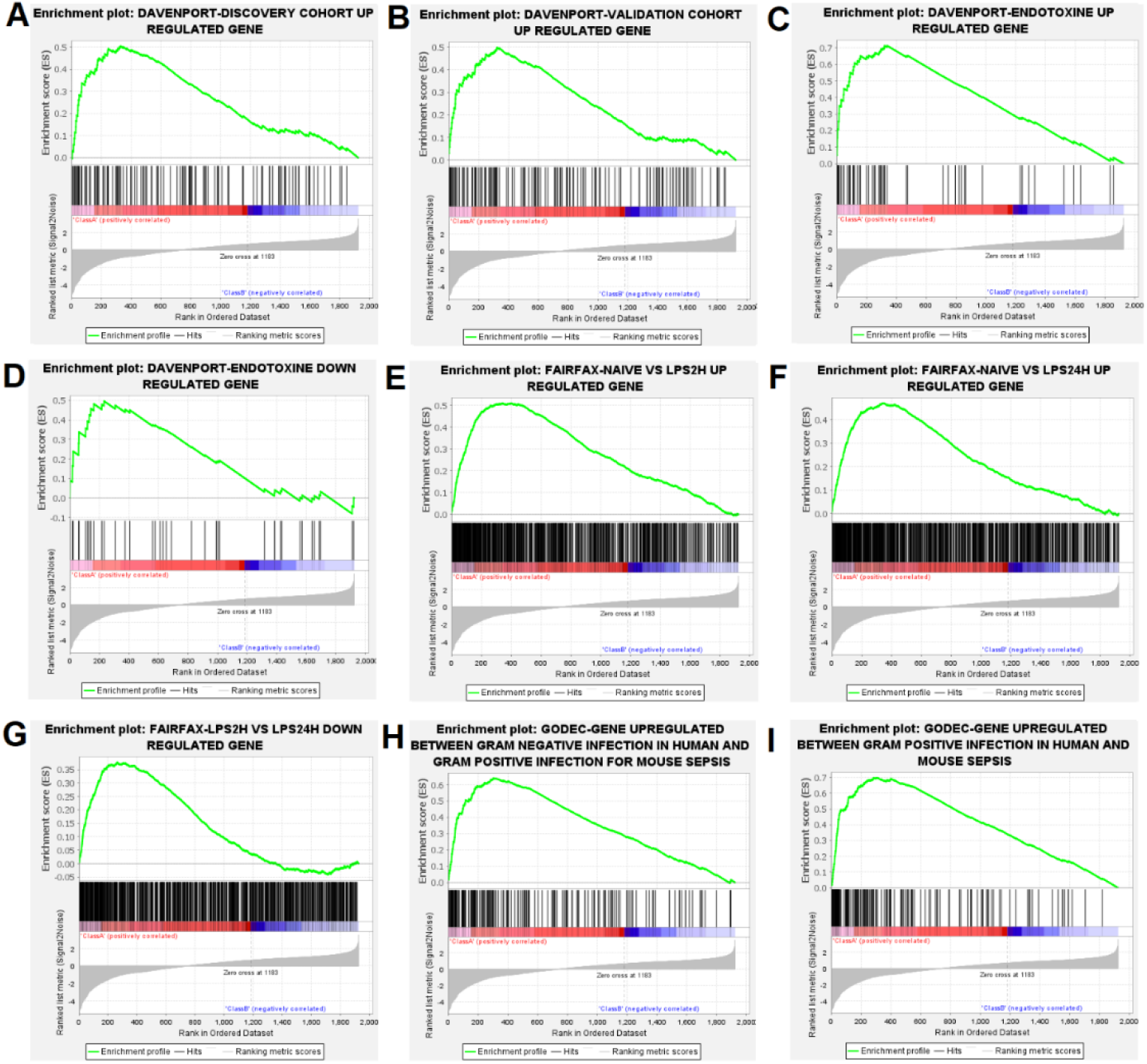
Gene set enrichment analysis (GSEA)-enrichment plots of comparison of our list of mouse differentially expressed genes with significant gene sets identified in 3 different papers. (A) Genes upregulated in the discovery cohort, (B) Genes upregulated in the validation cohort, (C) Genes upregulated in the endotoxin group (108 individuals from the discovery cohort with an immunosuppressed phenotype that included features of endotoxin tolerance), (D) Genes downregulated in the endotoxin group [8]. (E) Genes up-regulated in 2h LPS stimulated monocytes compared to naive monocytes, (F) Genes up-regulated in 24h LPS stimulate monocytes compared to naive monocytes, (G) Genes down regulated in 24h LPS stimulated monocytes compared to 2h LPS stimulated monocytes [5]. (H) Genes upregulated during GRAM negative infection in human and GRAM positive infection in mouse (GSE19668), (I) Genes up-regulated in GRAM positive infection in human (GSE9960) or mouse (GSE19668) [6].

**Figure 7:**
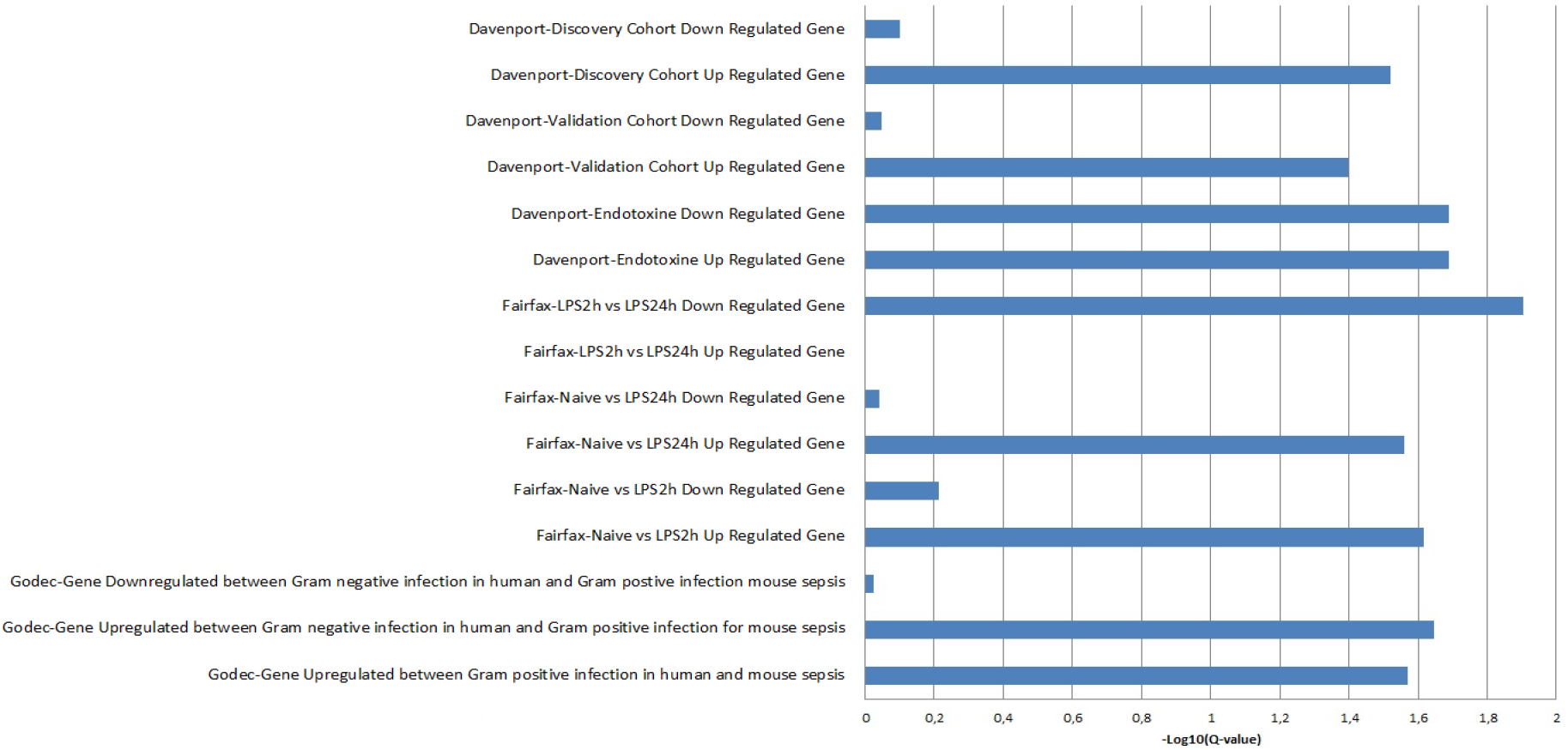
Plot of -Log10(Q-value) of gene set enrichment analysis for GSEA analysis.

In all, these results indicate that the response of mouse cells to LPS shares many features with that of human cells to LPS. This confirms previously reported results that the transcriptional response to sepsis is conserved in mouse and human peripheral blood mononuclear cells [6].

Also, we compared this published conserved gene expression profiles with the gene expression profile reported here: up-regulated genes in mouse cells stimulated with LPS for 1 hour were enriched in shared up-regulated genes which were previously published in human sepsis due to Gram-negative bacteria infection and in mouse sepsis due to Gram positive bacteria infection [6] (Figures 6H and 7). This supports the hypothesis that transcriptional host responses to Gram-positive bacteria and Gram-negative bacteria are very similar. In the same way, Tang *et al* reported that there were no signature genes that could differentiate between Gram-positive and Gram-negative sepsis in human peripheral blood mononuclear cells [7].

Moreover, up-regulated genes in mouse cells stimulated with LPS for 1 hour were enriched in human and mouse genes within gene sets associated with myeloid cells, whereas there was no enrichment in human and mouse genes within gene sets strongly associated with B or T lymphocytes [6] (Supplementary Figures 1 & 2).

Finally, we compared the list of the differentially expressed genes in LPS injected mice with human genes whose expression was associated with mortality in patients with sepsis [8]. GSEA approach yielded a significant enrichment after multiple test correction (Supplementary Table 3, Figure 6A, 6B, 6C, 6D, and Figure 7). Interestingly, Davenport et al compared their lists of genes with modulated genes in human peripheral mononuclear cells stimulated with LPS, as reported [9, 10]. They found 164 and 167 modulated genes in PBMC stimulated with LPS, which were up- and down regulated in susceptible patients from the discovery cohort, respectively. GSEA analysis yielded significant enrichment for up-regulated genes in the discovery and validation cohorts (Figures 6A, 6B and 7) and a significant enrichment for up and down regulated genes in the cohort of individuals with features of endotoxin tolerance (Figures 6C, 6D and 7). Among the 34 leading edge upregulated genes (in the group with endotoxin tolerance) identified by the GSEA method, we identified *IL1B, IL6, IL10* and *TNF* (Supplementary Table 3). Moreover, they were enriched in gene ontology terms related to inflammation, such as “inflammatory response”.

### 2.4. LincRNA study

The microarray contained 16,251 probes corresponding to lincRNAs and 4622 different lincRNAs. For each LincRNA, only the genomic position and the direction of transcription were filled in. For 12 of them, a gene symbol (BRN1-A, PINC, TUG1, HOTAIR, lincP21, NEAT2, NEAT1, NRON, H19, KCNQ1ot1, TSIX and XIST) was specified.

We looked then for a new and better annotation of the lincRNAs. For this purpose, we identify a symbol for the differentially expressed lincRNAs using their genomic positions in Gencode. Among the 126 differentially expressed lincRNAs in mice injected with LPS (Supplementary Table 1), we identified 1 lincRNA (Malat1). Malat1 (transcription of lung adenocarcinoma associated with metastasis) had a functional annotation and has been associated with lung cancers.

To better identify the putative role of lincRNA differentially expressed in our experiments, we looked for the role of neighboring coding genes and hypothesized that nearby lincRNA regulates the expression of this gene. To investigate this, we used the GREAT software (http://bejerano.stanford.edu/great/public/html/). Our lincRNAs list was submitted to recover the genes nearby (1000 kbp on each side of our lincRNA). The significant p-values <0.05 after the Benjamini correction were filtered, and the results are given in Supplementary Table 4. From 126 differentially expressed lincRNAs, approximately 208 genes were identified. Among those nearby genes, 40 coding genes were statistically differentially expressed in mice injected with LPS (Supplementary Table 4). Enrichment analysis results are shown in Supplementary Table 4 and concern a list of 208 genes associated with molecular functions such as the direct DNA binding, binding to regulatory regions of DNA, the downregulation of DNA-dependent transcription. We also noticed an enrichment for the Zinc finger proteins and motif prediction such as the GTF3A transcription factor. Of these, 21 clustered in the same group than the gene, suggesting a possible regulation of the gene regulation by the lincRNA (Supplementary Table 4). This is the case for BCL6 and Herc6 involved in the innate immune response in cluster 3 and 5, respectively and for ASXL1 and CRABP2 involved in the regulation of retinoic acid receptor signaling pathway in clusters 2 and 5, respectively.

## 3. Discussion

The relevance of the mouse model has been debated for several years [4, 11-14]. Seok et al. reported that transcriptional responses to acute inflammatory stresses in humans were poorly reproduced in the mouse models [15]. In contrast, Takao et al. detected a highly significant correlation between transcriptional responses in humans and those in mice using the same data sets and different analysis methods [16]. In the present study, we further investigated the genomic response to LPS in mice and compared it to that in humans.

We identified many differentially expressed genes in mouse peritoneal cells stimulated with LPS using an FDR median of 0%. Those genes were enriched in genes involved in inflammation, inflammatory disorders, immune suppression, antigen processing and presentation, apoptosis, and sepsis (pneumonia, sepsis, severe sepsis, and septic shock). Significant enrichments of differentially expressed genes were also identified in several signaling pathways, such as TNF-, IL-17, Chemokine-, NF-kappa B-, Toll-like receptor-, NOD-, MAPK-, JAK-STAT-signaling pathways. Interestingly, genes that were up-regulated in cells stimulated with LPS for 1 hour (Cluster 3) were enriched in genes involved in Toll-like receptor-, TNF-signaling, and NF-kappa B pathways, whereas other clusters did not show these enrichments. It should be noted that TNF and IL10 were up-regulated in cells stimulated with LPS for 1 hour, but not in cells stimulated with LPS for 40 hours. Furthermore, genes that were up-regulated in stimulated with LPS for 1 hour and 40 hours (cluster 6) were enriched in genes involved in response to interferon-gamma, chemokine signaling pathway, and regulation of chemotaxis for both natural killer cells and monocytes, whereas other clusters did not show these enrichments. Enrichment of IL-17- and JAK-STAT signaling pathways were evidenced for both clusters 3 and 6. It is likely that this reflects an early and transitional phase related to the production of TNF and a more progressive response related to the production of gamma IFN and chemokines. Noticeably, significant enrichment of immune suppression term was found in cluster 3, indicating that this process starts quickly after stimulation with LPS. Apoptosis of immune cells has been proposed to be a key mechanism for immune suppression in sepsis patients [17, 18]. Besides, IL-17 that has been shown to inhibit macrophage phagocytosis and to aggravate sepsis in a mouse model [19], likely participates in immune suppression.

In addition, our study of lincRNA located near differentially deregulated genes highlights the role of BCL6 known as an inhibitor of macrophage-mediated inflammatory responses [20-22] and involved in sepsis development as demonstrated in a recent publication in a mouse model [23].

It is very likely that macrophage peritoneal cells are mainly responsible for the response to LPS. Indeed, the molecular signature that yielded the best enrichment P value was that of peritoneal macrophage response to LPS for the whole list of differentially expressed genes, and for genes in cluster 3, 4, 5, and 6. This is also supported by the enrichment found for human monocytes stimulated by LPS, for other myeloid cells, but not for T and B lymphocytes. Also, our mouse model is restricted to a particular cell type. This is a limitation because other cell types, such as dendritic cells, T, B, and NK lymphocytes, and neutrophils are thought to be involved in the mechanisms of immune suppression [11, 24].

Genes up-regulated in mouse cells stimulated with LPS were enriched in genes up-regulated in human PBMC and monocytes stimulated with LPS [5, 8]. This confirms that the transcriptional response to LPS in mice reflects partly that of human cells. Moreover, genes up-regulated in mouse cells stimulated with LPS were enriched in genes up-regulated in susceptible sepsis patients [8]. Also, it is likely that there is a conserved sepsis molecular signature in human and mouse, as previously suggested by Godec et al. [6]. Noticeably, regulated genes in mouse cells stimulated by LPS were enriched in genes from the conserved sepsis molecular signature proposed by Godec et al.

These results show that mouse models can help to better understand the pathophysiology of sepsis in humans. In particular, mouse models can be used to establish a causal relationship between, on the one hand, death due to sepsis and, on the other hand, genes and molecular pathways, for which common features have been detected in humans and mice. Our results support that sepsis mouse model can be used to demonstrate working hypotheses for human sepsis. More particularly, one might take advantage of our mouse model to investigate the functional role of human candidate genes and pathways in monocytes or macrophages.

## 4. Materials and Methods

### 4.1. Mice and Experimental Procedure

Twelve C57BL/6SJL mice were purchased from the Charles River Laboratory (L’Arbresle, France). C57BL/6SJL mice were housed under specific pathogen-free conditions and handled in accordance with French and European directives (EU Agreement N° A-13013 03). To model severe form of sepsis, mice were intraperitoneally injected with 200 μL (35 mg/kg) of lipopolysaccharide from *Pseudomonas aeruginosa* (L9143, Sigma-Aldrich®). The mice were sacrificed by cervical rupture before injection T0 (n=4) and 1 hour after LPS injection during the inflammatory peak T1 (n=4) or 40 hours after LPS injection, just before the death T2 (n=4). Peritoneal cavity cells were collected by peritoneal washes using PBS.

### 4.2. RNA isolation and cDNA preparation for real-time RT-qPCR validation

Total RNA was extracted using the RNeasy extraction mini kit (QIAGEN, Hilden, Germany) and treated with DNase I (Qiagen). The quality of RNA was confirmed using an Agilent 2100 Bioanalyzer (Agilent Technologies, Germany) with Agilent RNA 6000 Nano Chips. Samples with a RIN higher than 8 were used. The quantity was measured using Qubit™ RNA Assay Kits on Qubit® 2.0 Fluorometer. For real-time RT-qPCR, reverse transcription reactions were performed using random hexamers and the SuperScript II reverse transcriptase (Invitrogen). Gene expression was evaluated using the SYBR green PCR master mix (PE Applied Biosystems, Foster, CA, USA) on Stratagene MX3000P (Agilent). Specific primers used are listed in Table 1.

**Table 1.**
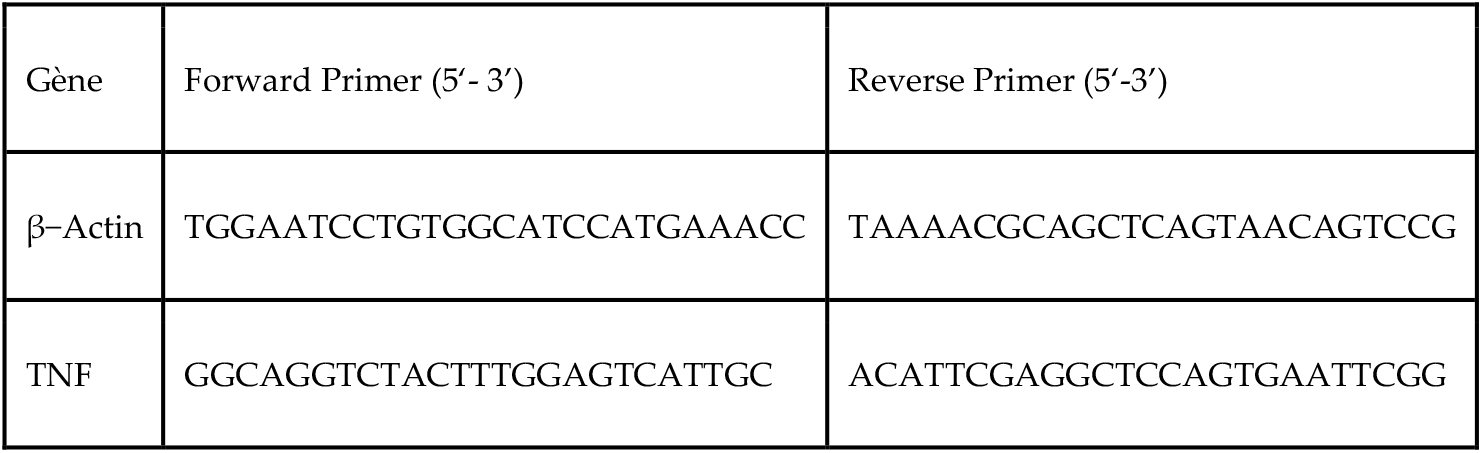
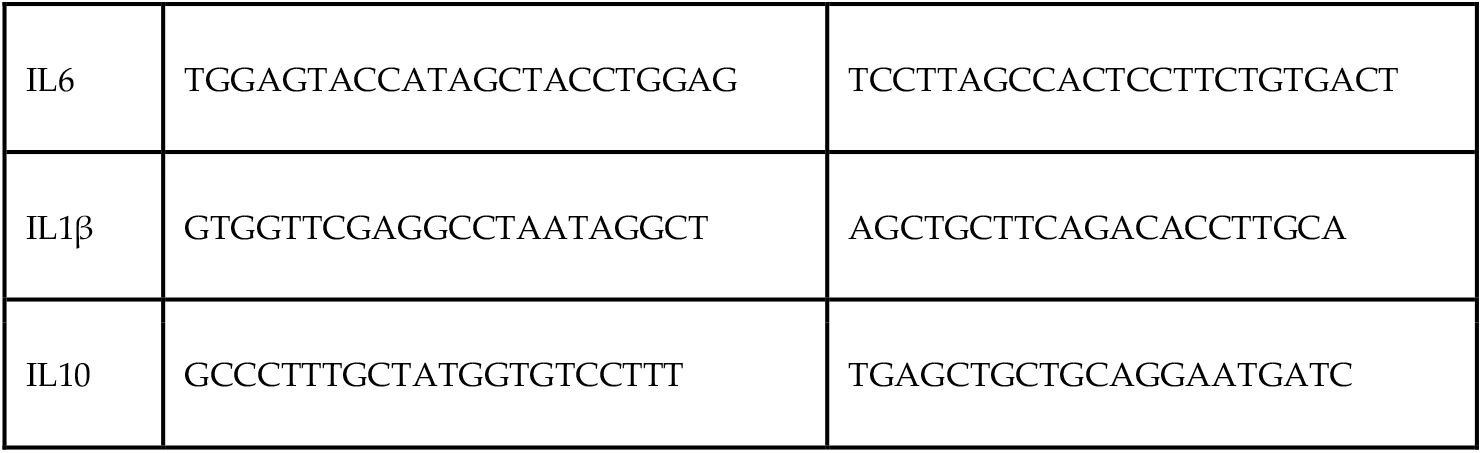
RT-qPCR primers sequences

### 4.3. Gene Expression Microarray

Labelling of RNA was done as recommended by Agilent Technologies using the One-Color Microarray-Based Gene Expression Analysis Low input Quick Amp Labeling. The Agilent microarray slides used is the SurePrint G3 mouse G3 8×60K chip harboring 62,976 probes including 39,430 coding RNAs and 16,251 lincRNAs (long intergenic nonRNA coding). The slides were scanned with “The Agilent Microarray Scanner®”. Visualization and results were retrieved using Feature Extraction® software. The raw data as well as the QC report were recorded. Twelve samples were used for analysis after quality validations.

### 4.4. Statistical analyses

The “AgiND” library loaded into R was used for standardizing microarray data. This library was developed for the diagnosis and standardization of Agilent chips. First, the raw data were transformed into log2 and then normalization was performed by the method of quantiles. A first filter was used to remove the controls, used to validate the correct hybridization. Then, a second filter was applied, allowing to remove the probes whose signal was close to the noise. Only 25880 probes with an expression greater than background noise were retained for all samples in a study group. For statistical analysis, we used TMeV software to perform a SAM (Significance Analysis of Microarray) to determine our differentially expressed genes. We used the False Discovery Rate (FDR) approach to correct for multiple tests; we applied an FDR median of 0%. Hierarchical clustering was further performed, using Pearson correlation coefficients, for the differentially expressed genes.

For RT-qPCR data, target gene levels were normalized based on B-actin level, which did not show any difference in microarray analysis, and quantified by the 2–ΔΔCt method [25] Kruskal-Wallis test was used to compare gene expression levels in different conditions. Spearman correlation test was performed to assess the correlation between microarray and RT-qPCR results for each gene.

### 4.5. Functional annotation and enrichment of functional terms

Functional analysis was performed with “Enrichr” to identify the biological processes and molecular function, in which genes are involved on the basis of “Gene Ontology” (GO) terms as well as the visualization of biological pathways from “KEGG” (Kyoto Encyclopedia of Genes and Genomes). We also checked the disease associated with these genes using DisGeNET, which is a discovery platform containing one of the largest publicly available collections of genes and variants associated with human diseases. The last database interrogated was Mouse Gene Atlas from BioGPS to find cell line enrichment in mouse tissues. The p-value was computed using a standard statistical method used by most enrichment analysis tools: Fisher’s exact test or the hypergeometric test. This is a binomial proportion test that assumes a binomial distribution and independence for the probability of any gene belonging to any set. Then, a p-value using the Benjamini-Hochberg method for correction for multiple hypotheses testing was applied to select the significant terms and pathways. In addition, an enrichment analysis was performed using the KEGG database to visualize the pathways of genes taking into account only the number of genes found. Q-values under 0.05 were considered significant. To assess the functionality of LincRNA impact by a bioinformatics approach, we used the Genomic Regions Enrichment of Annotation Tool (GREAT)[26]. GREAT evaluates sets of Cis-regulatory elements by assigning each element to its likely target gene(s).

### 4.6. Comparison of published lists of human differentially expressed genes with mouse differentially expressed genes

Our mouse gene expression analysis was compared to human datasets published by Fairfax et al [5] and Davenport et al [8] Gene sets published were directly used in our analysis. Table 2 summarizes the characteristics of the studies. In addition, our mouse gene expression analysis was compared to conserved patterns of gene expression in humans and mouse models triggered by sepsis [6].

**Table 2.** Characteristics of human datasets in the articles of Davenport et al[8] and Fairfax et al[5]

|  | Davenport-Discovery cohort | Davenport-Validation cohort | Fairfax-Naive vs 2h LPS<br>stimulated<br>monocytes | Fairfax-Naive vs 24h LPS<br>stimulated<br>monocytes | Fairfax-2h LPS vs 24h<br>LPS stimulated<br>monocytes |
| --- | --- | --- | --- | --- | --- |
| **Chip/Paper** | Illumina Human-HT-12 version 4<br>Expression<br>BeadChips with<br>47 231 probes | Illumina Human-HT-12 version 4<br>Expression<br>BeadChips with<br>47 231 probes | Illumina<br>HumanHT-12 v4<br>BeadChip gene<br>expression array<br>platform with<br>47,231 probes | Illumina<br>HumanHT-12 v4<br>BeadChip gene<br>expression array<br>platform with<br>47,231 probes | Illumina<br>HumanHT-12 v4<br>BeadChip gene<br>expression array<br>platform with<br>47,231 probes |
| **Number of**<br>**samples** | 270 patients with<br>sepsis due to<br>pneumonia and<br>organ dysfunction | 114 patients with<br>sepsis due to<br>pneumonia and<br>organ dysfunction | 228 individuals /<br>LPS (E.Coli) for 2h | 228 individuals /<br>LPS (E.Coli) for<br>24h | 228 individuals /<br>LPS (E.Coli) for<br>2h or 24h |
| **Statistic** | linear regression<br>using limma | linear regression<br>using limma | linear regression<br>using limma | linear regression<br>using limma | linear regression<br>using limma |
| **Cut-off** | false discovery<br>rate of 0.05 and<br>1.5-fold change in<br>expression<br>between the two<br>groups | false discovery<br>rate of 0.05 and<br>1.5-fold change in<br>expression<br>between the two<br>groups | Adjusted pvalue<br>and cut-off fold<br>change of +/-0.5 | Adjusted pvalue<br>and<br>cut-off fold<br>change of +/-0.5 | Adjusted pvalue<br>and cut-off fold<br>change of +/-0.5 |
| **Gene**<br>**Differentially**<br>**expressed** | 3080 (331) | 2572 | 2489 | 2085 | 2913 |
| Up-regulated genes | 821 (164) | 1117 | 1564 | 1116 | 1169 |
| Down-regulated genes | 2260 (167) | 1463 | 925 | 969 | 1744 |
| Common Up/Down | 1 (0) | 8 | 0 | 0 | 0 |

Gene Set Enrichment Analysis (GSEA) was performed using GSEA software [27]. Given the gene sets previously published, we assessed whether the members of those gene sets were randomly distributed throughout the list of mouse differentially expressed genes, which were ranked on the basis of the SAM score. Enrichment scores and the corresponding nominal P values were calculated. Multiple testing was controlled by calculating the FDR. An FDR under 5% was considered significant.

## Supplementary Materials

The following are available online at www.mdpi.com/xxx/s1, Figure S1: title, Table S1: title, Video S1: title.

## Author Contributions

F.R. performed the experiments with mice, extracted RNA, prepared cDNA and performed the qRT-PCR; he analysed the results using bioinformatic tools; he participated in writing the paper. N.F.N. and B.L. performed the Gene Expression Microarray. M.T. participated in the experiment with mice and qRT-PCR. P.R. co-supervised the statistical analyses, participated extensively in the paper writing. L.C.P. performed the experiment with mice, co-supervised the experimental studies and participated extensively in the paper writing. P.R. and L.C.P. jointly supervised this work.

## Funding

This work was supported by the French Research Ministry, by the Institut National de la Santé et de la Recherche Médicale (INSERM). This work has not been presented previously at a scientific meeting.

## Institutional Review Board Statement

Mice used in this study were housed and treated according to the conditions of the Platform for Stabling and Animal Experimentation (PSEA, Parc Scientifique de Luminy, 13288 Marseille cedex 9, France) which is a member of the national network CELPHEDIA (WP Zootechnics, Ethics & Animal Welfare group). The project involving the mice was submitted to the ethics committee for animal experimentation under the number 202110161713613.

## Data Availability Statement

The transcriptomic data are available in the GEO database (GSE185150). The data used to support the findings of the study are available from the corresponding author upon request.

## Acknowledgments

We thank the TGML Platform, supported by grants from Inserm, GIS IBiSA, Aix-Marseille Université, and ANR-10-INBS-0009-10. We thank Aurélie Bergon for depositing the transcriptomic data in the GEO database.

## Conflicts of Interest

The authors declare that there is no conflict of interests regarding the publication of this article.

## Notes

### Competing Interest Statement

The authors have declared no competing interest.

