## Supplementary figures and images for "Transcriptional response in a sepsis mouse model reflects transcriptional response in sepsis patients"

### Supplementary Figure 1.tiff

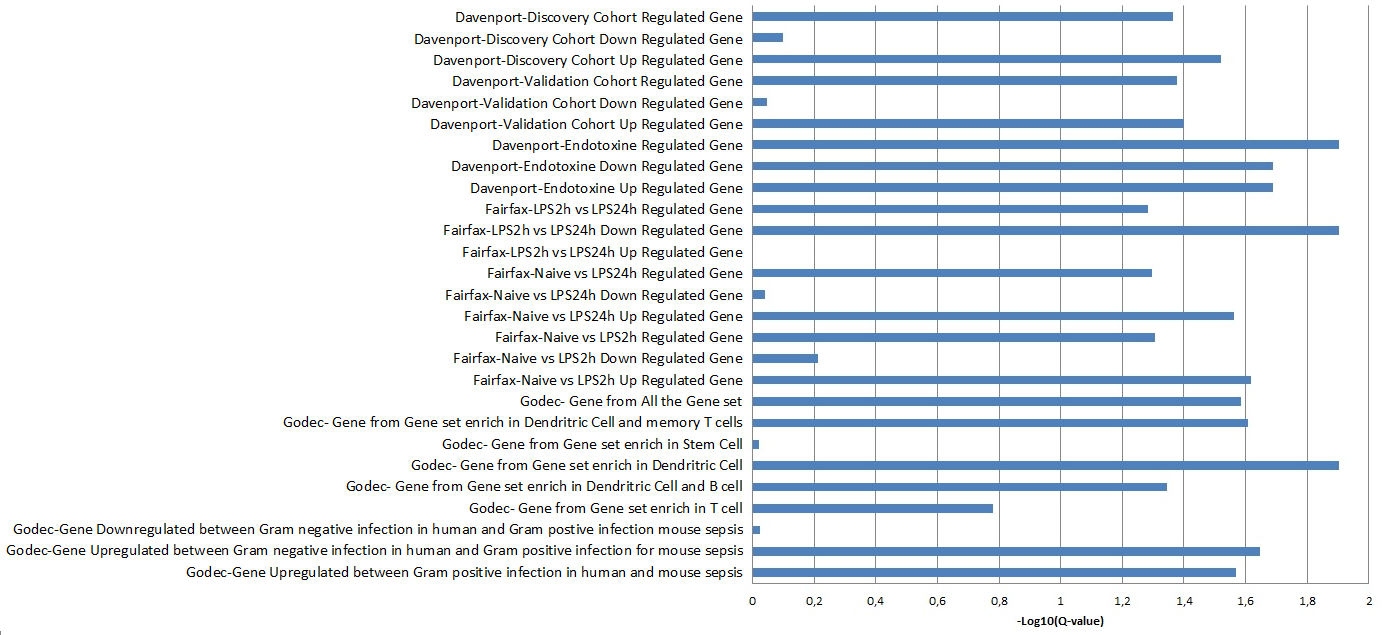

### Supplementary Figure 2.tiff

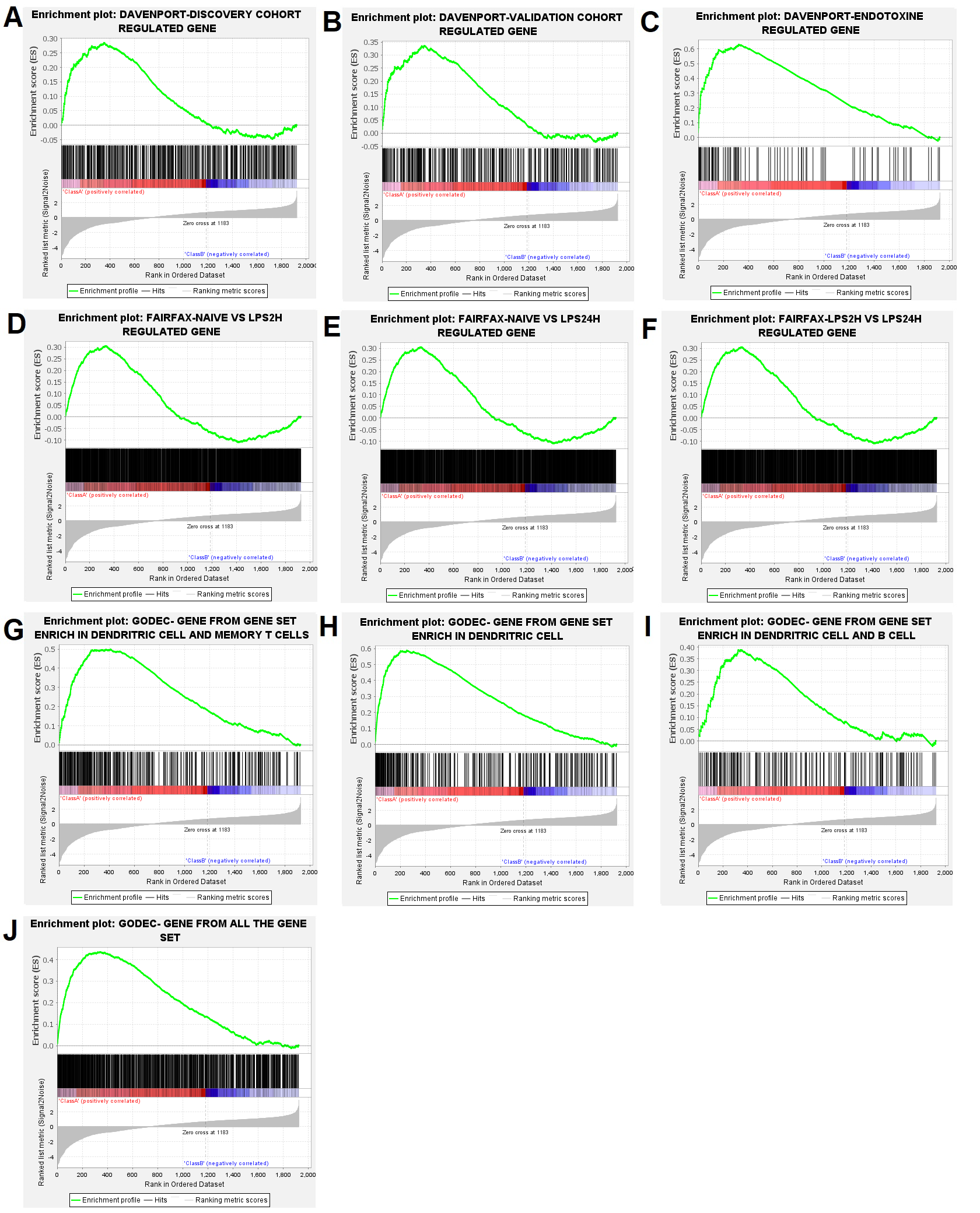
